# Extracellular CIRP as a Novel Endogenous TREM-1 Ligand to Fuel Inflammation

**DOI:** 10.1101/674218

**Authors:** Naomi-Liza Denning, Monowar Aziz, Atsushi Murao, Steven D. Gurien, Mahendar Ochani, Jose M. Prince, Ping Wang

## Abstract

Extracellular cold-inducible RNA-binding protein (eCIRP) is a recently-discovered damage-associated molecular pattern. Understanding the precise mechanism by which it exacerbates inflammation in sepsis is essential. Here we identified that eCIRP is a new biologically active endogenous ligand of triggering receptor expressed on myeloid cells-1 (TREM-1), fueling inflammation in sepsis and ischemia-reperfusion. Surface plasmon resonance revealed a strong binding affinity between eCIRP and TREM-1, and FRET assay confirmed eCIRP’s interaction with TREM-1 in macrophages. TREM-1 inhibition, either by its siRNA or a decoy peptide LP17, dramatically reduced eCIRP-induced inflammation. We developed a novel 7-aa peptide derived from human eCIRP, M3, which blocked the interaction of TREM-1 and eCIRP. M3 suppressed inflammation induced by eCIRP or agonist TREM-1 Ab crosslinking in murine macrophages or human peripheral blood monocytes. M3 also inhibited eCIRP-induced systemic inflammation and tissue injury. Treatment with M3 further protected mice from sepsis and intestinal ischemia-reperfusion, improved acute lung injury, and increased survival. Thus, we have discovered a novel TREM-1 ligand and developed a new peptide M3 to block eCIRP-TREM-1 interaction and improve the outcome of sepsis and sterile inflammation.

## Introduction

Pathogen-associated molecular patterns (PAMPs) and damage-associated molecular patterns (DAMPs) play leading roles in fueling inflammation in both infectious and sterile systemic insults, such as those that occur during sepsis or ischemia/reperfusion (I/R) injury ^1, 2, 3^. Cold-inducible RNA-binding protein (CIRP) is a 18-kDa RNA chaperone protein ^4^. In addition to the passive release due to necrotic cell death, CIRP can be released extracellularly during sepsis, hemorrhage or I/R injury, translocating from the nucleus to cytoplasmic stress granules before being released to the extracellular space ^5^. Extracellular CIRP (eCIRP) acts as a DAMP, causing severe inflammation and organ injury ^5, 6^. Elevated plasma levels of eCIRP have been correlated with a poor prognosis in patients with sepsis ^5, 7^. Being a new DAMP, a complete understating of eCIRP’s pathobiology in inflammatory diseases is required to develop novel therapeutics.

Similar to other DAMPs, eCIRP recognizes Toll-like receptor 4 (TLR4) ^5, 8^ expressed in macrophages, lymphocytes, and neutrophils to increase cytokine production ^5^, T cell polarization ^9^, and neutrophil activation ^10^. Healthy mice injected with recombinant murine (rm) CIRP developed acute lung injury (ALI) via activation of macrophages, neutrophils, and endothelial cells, and displayed increased vascular permeability in the lungs ^11, 12^. Conversely, CIRP^-/-^ mice were protected from sepsis and ALI ^5, 6^. Consistent with these findings, treatment of septic animals with a neutralizing antibody against eCIRP attenuated organ injury and prolonged survival ^5, 6^. Collectively these findings indicate that eCIRP is a major contributing factor in the pathogenesis of sepsis; targeting eCIRP is a valid strategy to mitigate sepsis severity.

Triggering receptor expressed on myeloid cells-1 (TREM-1) is an innate immune receptor expressed primarily on neutrophils and macrophages ^13^. TREM-1 activation triggers inflammation independently ^14^, as well as by synergizing with the TLR4 pathways ^14, 15^. TREM-1 activation leads to DNAX-activating protein of 12 kDa (DAP12) phosphorylation ^13, 15^ which promotes activation of the tyrosine kinase Syk ^16^, resulting in the production of cytokines and chemokines ^15, 16^. The ligands for TREM-1 remain elusive, with only high mobility group box 1 (HMGB1) ^17^, extracellular actin ^18^, and peptidoglycan recognition protein 1 (PGLYRP1) ^19^ identified thus far. The nature of the TREM-1 ligand(s) and mechanisms of TREM-1 signaling are not yet thoroughly explored. Identification of a new natural endogenous TREM-1 ligand will not only help improve our understanding of the pathophysiology of inflammatory diseases, but also discover new therapeutic avenues against those diseases.

Both eCIRP and TREM-1 are upregulated in sepsis to serve as mediators of inflammation ^5, 20^, but their interaction has not been studied. In sepsis, both eCIRP and TREM-1 reside extracellularly ^5, 20^, giving rise to the hypothesis that eCIRP could be a novel endogenous ligand of TREM-1. In this study, we have discovered that eCIRP is a new biologically active endogenous TREM-1 ligand and their interaction fuels inflammation. We have also developed a unique human eCIRP-derived ligand-dependent 7 amino acid peptide (RGFFRGG) to serve as an antagonist of TREM-1, named M3, and implemented M3 as a therapeutic in pre-clinical models of sepsis and I/R injury. Our findings provide an effective therapeutic target against sepsis and sterile inflammatory diseases.

## Results

### Identification of eCIRP as a new TREM-1 ligand to promote inflammation

To study the direct interaction between eCIRP and TREM-1, we performed a surface plasmon resonance (SPR) assay, which demonstrated a strong binding between rmCIRP and rmTREM-1 with a KD of 11.7 × 10^-8^ M (Fig 1A). An immunofluorescence study was performed to confirm the co-localization of eCIRP and TREM-1 in macrophages after rmCIRP stimulation. It clearly demonstrated the co-localization of rmCIRP and TREM1, as indicated by the merged (yellow) image (Fig 1B). Conversely, rmCIRP did not co-localize with a negative control, the macrophage pan marker CD11b (Fig 1B). We next performed FRET analysis to quantitatively determine rmCIRP’s association with TREM-1. FRET analysis revealed a clear association between rmCIRP and TREM-1 with an increase in FRET units of nearly 7-fold compared to rmCIRP’s interaction with negative control CD11b (Fig 1C). These findings implicate that eCIRP is a novel TREM-1 ligand. We then studied the activation of downstream molecules DAP12 and Syk in macrophages treated with rmCIRP and found robust increase in the phosphorylation of DAP12 and Syk at 10 min after rmCIRP stimulation (Fig 1D-F; **Supplemental Fig 1A-E**). We next confirmed the functional role of TREM-1 in eCIRP-mediated inflammation. We found that the siRNA-treated macrophages showed significant inhibition of TNF-α production following rmCIRP stimulation (Fig 1G). Similarly, the treatment of macrophages with LP17, an inhibitor of TREM-1 ^21^, dose-dependently inhibited rmCIRP-induced TNF-α production in RAW264.7 cells (Fig 1H). Conversely, the scramble peptide for LP17 did not show any inhibition of TNF-α production (Fig 1H). Collectively, these data clearly show that eCIRP specifically binds to TREM-1 in macrophages and induces TNF-α production. TREM-1 expression in macrophages is increased in sepsis ^15^. To explore the role of eCIRP on this increase, RAW264.7 cells and murine primary peritoneal macrophages were stimulated with rmCIRP. TREM-1 mRNA levels were increased 2.5-fold in rmCIRP-treated RAW264.7 cells as compared to PBS control **(Supplemental Fig 2A)**. The protein levels of TREM-1 expression on the cell surface of both RAW264.7 cells and primary murine peritoneal macrophages treated with rmCIRP were significantly increased by 4.3 and 1.6-fold, respectively, compared to PBS control **(Supplemental Fig 2B, C).**

**Figure 1:**
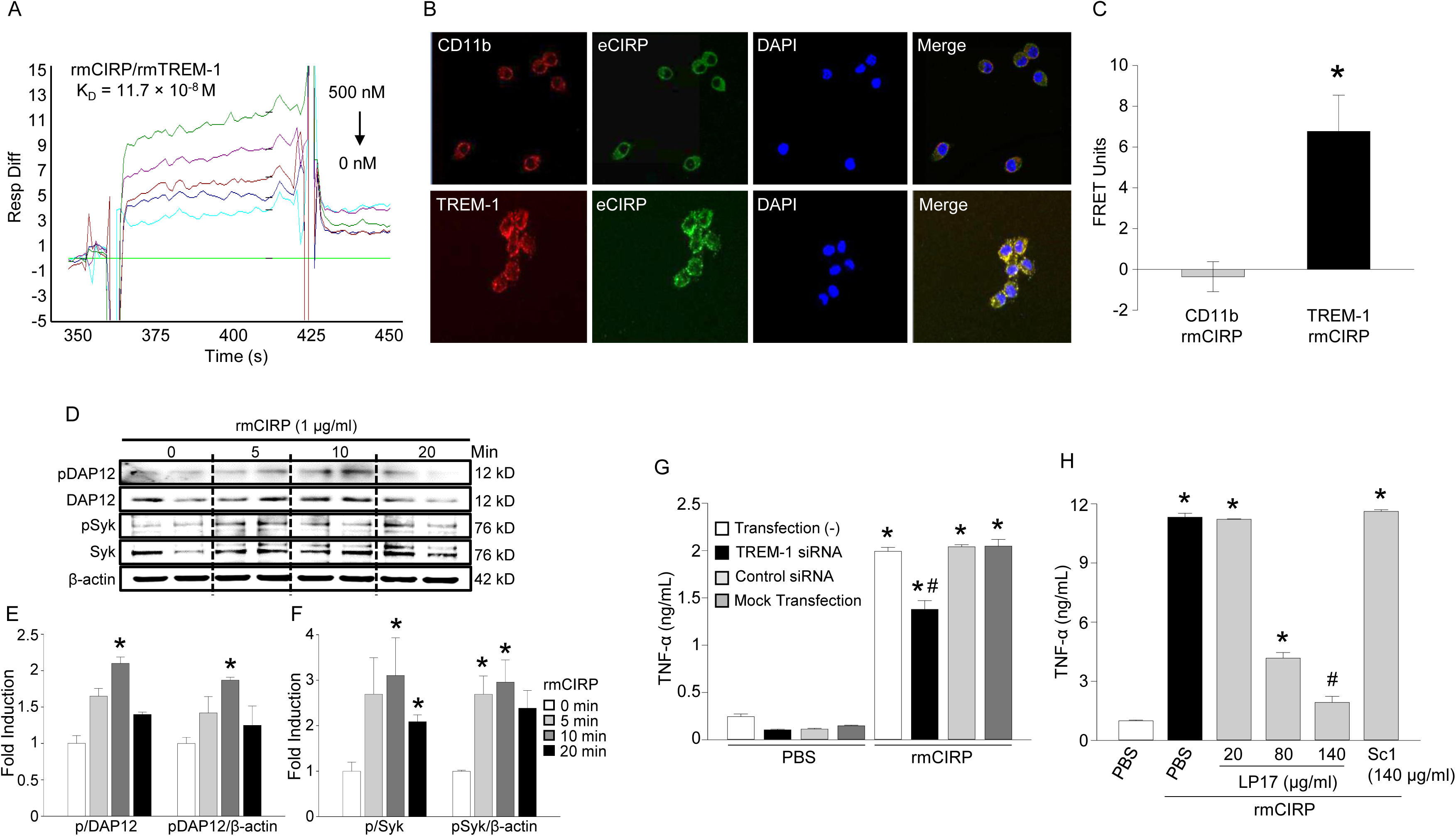
eCIRP binds TREM-1 to promote inflammation. **(A)** SPR between rmCIRP and rmTREM-1. Anti-his Ab was used to capture rmCIRP-his. rmTREM-1 was injected as an analyte in concentrations of 0 to 500 nM. **(B)** 1.5 × 10^4^ RAW264.7 cells were treated with rmCIRP (5 μg/ml) at 4°C for 10 min, fixed in a nonpermeabilized fashion, and stained with rabbit anti-mouse CIRP Ab, goat anti-mouse TREM-1 Ab, goat anti-CD11b Ab, Cy5-conjugated AffiniPure donkey anti-goat IgG, and Cy3-conjugated AffiniPure F(ab’)2 fragment donkey anti-rabbit IgG. Confocal microscopy images were obtained at using a Zeiss LSM880 confocal microscope equipped with a 63× objective. Images were analyzed and quantified by using the ZenBlue software. (**C)** After the staining protocol described in **(B)**, cell associated fluorescence was measured on Biotek Synergy Neo2 at 566 nm upon excitation at 488 nm (*E*1), at 681 nm after excitation at 630 nm (*E*2), and at 681 nm after excitation at 488 nm (*E*3). The transfer of fluorescence was calculated as FRET units. FRET unit = [*E*3_both_ −*E*3_none_] − [(*E*3_Cy5_ −*E*3_none_) × (*E*2_both_/*E*2_Cy5_)] − [(*E*3_Cy3_ − *E*3_none_) × (*E*1_both_/*E*1_Cy3_)]. Data are expressed as means ± SE obtained from three independent experiments, n=9/group. Groups compared by unpaired t-test (*p<0.01 vs. CD11b). **(D)** A total of 1 × 10^6^/ml RAW264.7 cells were stimulated with rmCIRP (1 µg/ml) for various times. Extracted proteins were immunoprecipitated by using anti-DAP12 Ab, followed by Western blotting using pTyr (4G10) and DAP12 Ab. Extracted proteins obtained from rmCIRP (1 µg/ml for various times) stimulated RAW264.7 cells (1 × 10^6^/ml) were subjected to Western blotting using pSyk, Syk, and β–actin Abs. Representative western blots for phosphotyrosine (4G10), DAP12, pSyk, Syk, and β–actin are shown. **(E, F)** Each blot was quantified by densitometric analysis. Phosphotyrosine (pDAP12) and pSyk expression in each sample was normalized to DAP12 or Syk or β-actin expression and the mean values of 0 min of rmCIRP-treated groups were standardized as one for comparison. Data are expressed as means ± SE (n=6 samples/group). The groups were compared by one-way ANOVA and SNK method (*p<0.05 vs. rmCIRP at 0 min). **(G)** RAW264.7 cells were transfected with TREM-1 siRNA, control siRNA, underwent mock transfection, or no transfection. Cells were then stimulated with PBS control or 1μg/ml rmCIRP. After 6 h, TNF-α in the supernatant was analyzed by ELISA. Data are expressed as means ± SE (n=3 samples/group). Multiple groups were compared by one-way ANOVA and Tukey method (*p<0.05 vs. respective PBS group; ^#^p<0.01 vs. rmCIRP-treated non-transfected cells). **(H)** RAW264.7 cells (1×10^4^ cells/ml) were plated in 96-well culture plate and stimulated with PBS or rmCIRP (1 µg/ml). Simultaneously cells were treated with various doses of LP-17 or LP-17-Sc1. After 24 h, TNF-α in culture supernatants at protein level were measured by ELISA. Data are expressed as means ± SE obtained from five independent experiments (n=3-10 wells/group). The groups were compared by one-way ANOVA and SNK method (*p<0.05 vs. PBS; #p<0.05 vs. rmCIRP+PBS). FRET, fluorescence resonance energy transfer; TNF, tumor necrosis factor; DAP12, DNAX activation protein of 12kDa; ELISA, enzyme-linked immunosorbent assay; PBS phosphate buffered saline.

### An eCIRP-derived antagonist M3 inhibits the eCIRP-TREM-1 interaction

After establishing eCIRP as a novel TREM-1 ligand, we sought to identify a small molecule that inhibits the eCIRP-TREM-1 interaction. Employing the Protein Model Portal, part of the protein structure initiative knowledgebase, and Pep-Fold-3 ^22^, we compared structural images of a known TREM-1 ligand, murine PGLYRP1, and murine CIRP. We identified a section of CIRP with a similar structural form and associated similar amino acid sequences (**Supplemental Fig 3A-B**). We then synthesized a series of small peptides (M1, M2, M3) from this region of CIRP and tested their ability to inhibit TNF-α secretion in RAW264.7 cells stimulated with rmCIRP (Fig 2A). We found that M3, a 7-aa peptide, consisting of the amino acid sequence of murine CIRP from 101-107 (RGFFRGG) which has 100% homology with human CIRP demonstrated the greatest inhibitory effect on TNF-α production by the macrophages following rmCIRP treatment (Fig 2A). Using SPR, we identified considerable binding affinity between M3 and rmTREM-1 with a KD of 35.2 × 10^-6^ M (Fig 2B). FRET assay was performed between rmCIRP and TREM-1 in macrophages in presence or absence of M3. Interestingly, we found that M3 was able to dramatically abrogate rmCIRP’s binding to TREM-1 (Fig 2C). The agonist anti-TREM-1 Ab has been shown to induce TNF-α production through TREM-1 crosslinking ^13^. By inhibiting TREM-1-mediated inflammation caused solely by antibody induced receptor activation in the absence of any other stimuli, we have demonstrated the independent role of M3 on TREM-1-mediated inflammation. We found that M3 demonstrated significant inhibition (61%) of TNF-α production by the macrophages treated with an agonist anti-TREM-1 Ab, indicating M3 specifically worked on TREM-1 (Fig 2D). On the other hand, the scramble peptides, M3-Sc1 and M3-Sc2 did not demonstrate any inhibition (Fig 2D). We next validated M3’s inhibitory effect on controlling rmCIRP-treated inflammation in macrophages. We found that M3 inhibited rmCIRP-mediated TNF-α production by the RAW264.7 cells in a dose-dependent manner while the scramble peptides M3-Sc1 and M3-Sc2 were not able to inhibit rmCIRP-induced TNF-α production (Fig 2E). Similarly, M3 treatment was also able to significantly inhibit the production TNF-α in rmCIRP-stimulated primary human PBMC obtained from healthy volunteers where the highest inhibition of 76% in TNF-α production occurred at 10 μg/ml of M3 treatment (Fig 2F). Collectively, we have discovered a novel small peptide M3 which blocks the eCIRP-TREM-1 interaction and inhibits eCIRP-mediated inflammation.

**Figure 2:**
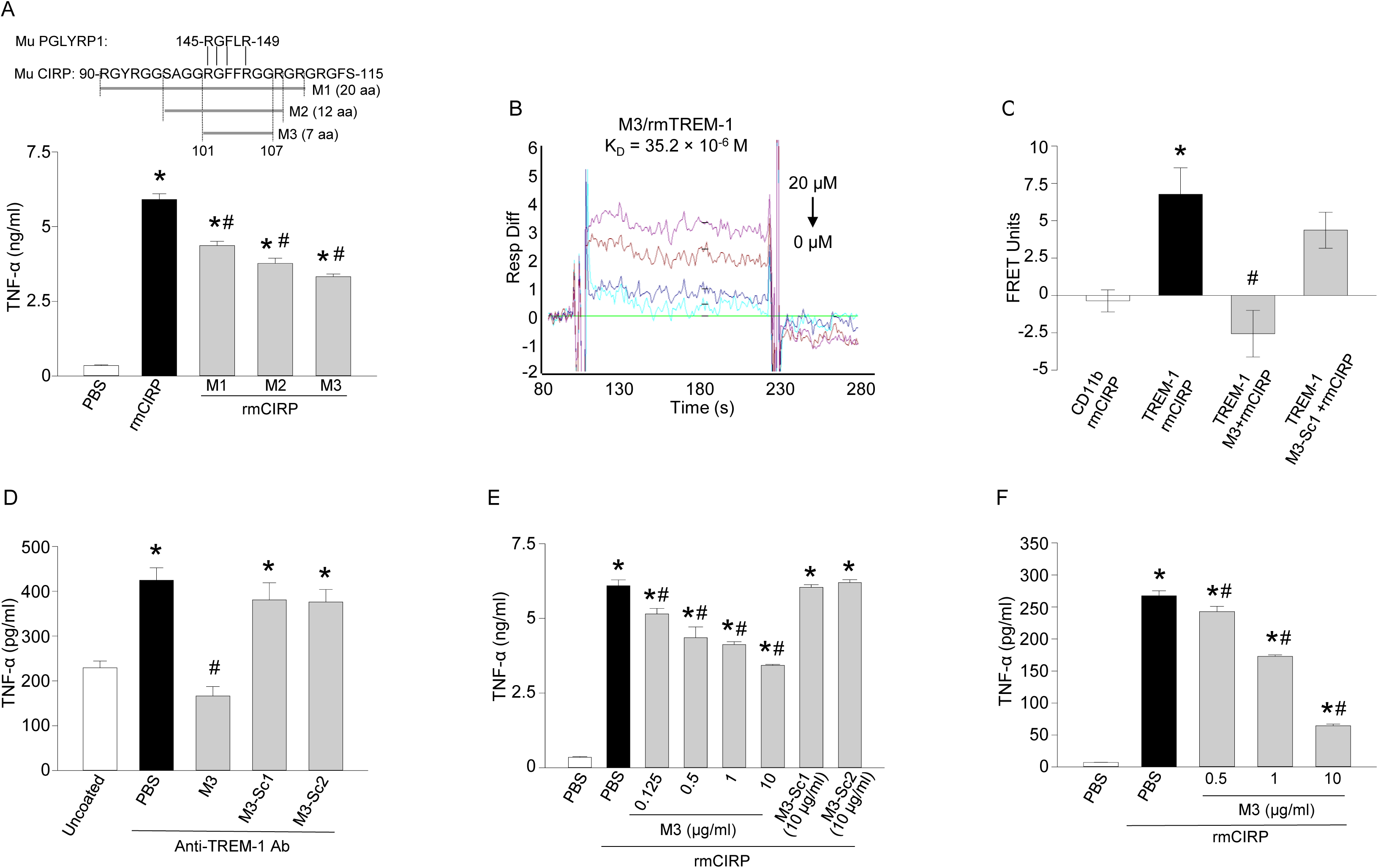
M3, a small CIRP-derived peptide, inhibits eCIRP and TREM-1 interaction. **(A)** Partial amino acid sequence of murine CIRP highlighting an area of similarity between murine PGLYRP1. Three peptides (M1, M2, and M3) are highlighted from within the CIRP sequence. RAW264.7 cells (1×10^4^ cells/ml) were plated in 96-well culture plate and treated with 10 µg/ml of peptides M1, M2, or M3 for 30 min. Cells were then stimulated with rmCIRP (1 µg/ml). After 24 h, TNF-α in culture supernatants at protein level were measured by ELISA. Data are expressed as means ± SE obtained from two independent experiments (n=6 wells/group). The groups were compared by one-way ANOVA and SNK method (*p<0.05 vs. unstimulated cells and #p<0.05 vs. rmCIRP-treated cells). **(B)** SPR between rmTREM-1 and M3. M3 was injected as an analyte in concentrations of 0 to 20 μM. **(C)** RAW264.7 cells (1×10^4^ cells/ml) were plated in 96-well culture plate and treated with M3 or M3-Sc1 at a dose of 10 µg/ml for 30 min. Cells were then stimulated with PBS or rmCIRP (5 µg/ml) at 4°C for 10 min, fixed in a nonpermeabilized fashion, and stained as described in 1B. FRET analysis was performed as described in 1B. Data are expressed as means ± SE obtained from three independent experiments, n=9/group. Multiple groups were compared by one-way ANOVA and Tukey method (*p<0.05 vs CD11b + rmCIRP, #p<0.05 vs TREM-1 + rmCIRP). **(D)** To activate RAW264.7 cells through TREM-1, 96-well flat bottom plates were pre-coated with 20 μg/ml of an agonist anti-TREM-1 mAb overnight at 37°C. The wells were washed with sterile PBS and 5 × 10^4^ cells/well were plated. Prior to plating, cells were premixed with either PBS control, M3 (10 μg/ml) or scramble M3-Sc1 (10 μg/ml) for 30 min. After plating, TNF-α production was measured in the culture supernatants after an additional 24 h of incubation. Data are expressed as means ± SE. The experiment was performed two independent times with n=5 wells per group. Multiple groups were compared by one-way ANOVA and Tukey method (*p<0.05 vs. uncoated; #p<0.05 vs. TREM-1 Ab + PBS). **(E)** RAW264.7 cells (1×10^4^ cells/ml) were plated in 96-well culture plate and treated with various doses of M3, M3-Sc1, or M3-Sc1 for 30 min. Cells were then stimulated with PBS or rmCIRP (1 µg/ml). After 24 h, TNF-α in culture supernatants at protein level were measured by ELISA. Data are expressed as means ± SE obtained from three independent experiments (n=4 wells/group). The groups were compared by one-way ANOVA and SNK method (*p<0.05 vs. PBS-treated cells; #p<0.05 vs. rmCIRP + PBS). **(F)** Macrophages from healthy human donors (1×10^4^ cells/ml) were plated in 96-well culture plate and treated with various doses of M3 for 30 minutes. Cells were then stimulated with PBS or rmCIRP (1 µg/ml). After 24 h, TNF-α in culture supernatants at protein level were measured by ELISA. Data are expressed as means ± SE (n=5 wells/group). The groups were compared by one-way ANOVA and SNK method (*p<0.05 vs. PBS-treated cells; #p<0.05 vs. rmCIRP + PBS). FRET, fluorescence resonance energy transfer; TNF, tumor necrosis factor; ELISA, enzyme-linked immunosorbent assay; PBS, phosphate buffered saline; PGLYRP1, peptidoglycan recognition protein 1.

### M3 inhibits eCIRP or LPS-induced inflammation in mice

Having established M3 as an antagonist of eCIRP and TREM-1 binding, we sought to evaluate its efficacy in reducing systemic inflammation *in vivo*. Administration of rmCIRP in healthy mice dramatically increased serum levels of IL-6 and IL-1β, and M3 treatment decreased these levels by 50% and 24%, respectively (Fig 3A, B). Lung mRNA levels of TNF-α, IL-1β, and KC were significantly increased by rmCIRP injection into mice, while M3 treatment reduced their expression by 65%, 74%, and 45%, respectively (Fig 3C-E). We further proved that pharmacologic inhibitor of TREM-1 using LP17 attenuated rmCIRP-induced systemic inflammation and lung injury in mice (Fig 4). Treatment of mice with LP17 dramatically reduced the levels of AST, ALT, and LDH in the serum of rmCIRP-injected mice by 77%, 82%, and 54%, respectively, compared to vehicle-treated rmCIRP-injected mice (Fig 4A-C). The levels of IL-6, IL-1β, and IFN-γ in the serum were significantly decreased by 72%, 66%, and 86%, respectively in LP17-treated mice (Fig 4D-F). LP17 treatment significantly attenuated TNF-α, IL-1β, and IL-6 mRNA by 80%, 59%, and 93%, respectively, (Fig 4G-I) and proteins by 24%, 63%, and 30% respectively, in the lungs (Fig 4J-L). Similarly, treatment with LP17 significantly reduced the expression of chemokines MIP-2 and KC and adhesion molecules ICAM-1 and VCAM-1 mRNA in the lungs (**Supplemental Fig 4A-D**). Histological images of lung tissue showed increased levels of alveolar congestion, exudate, interstitial and alveolar cellular infiltrates, intra-alveolar capillary hemorrhages, and damage of epithelial architecture, in rmCIRP-injected mice compared to sham mice (Fig 4M). LP17 treatment dramatically improved these histological injury parameters in rmCIRP-injected mice (Fig 4M). These histological changes were reflected in a significant decrease in lung tissue injury score in LP17-treated mice by 48% (Fig 4N). LPS is known to stimulate eCIRP release by the macrophages ^5^. Using an endotoxemia model, we demonstrated that M3 treatment was able to reduce serum levels of IL-6 and TNF-α by 31% and 80%, respectively (Fig 3F, G). Administration of M3 during endotoxemia increased the 7-day survival from 70% to 100% (Fig 3H). Therefore, TREM-1 is involved in eCIRP-mediated inflammation and tissue injury, and blockade of the eCIRP-TREM-1 interaction attenuates inflammation.

**Figure 3:**
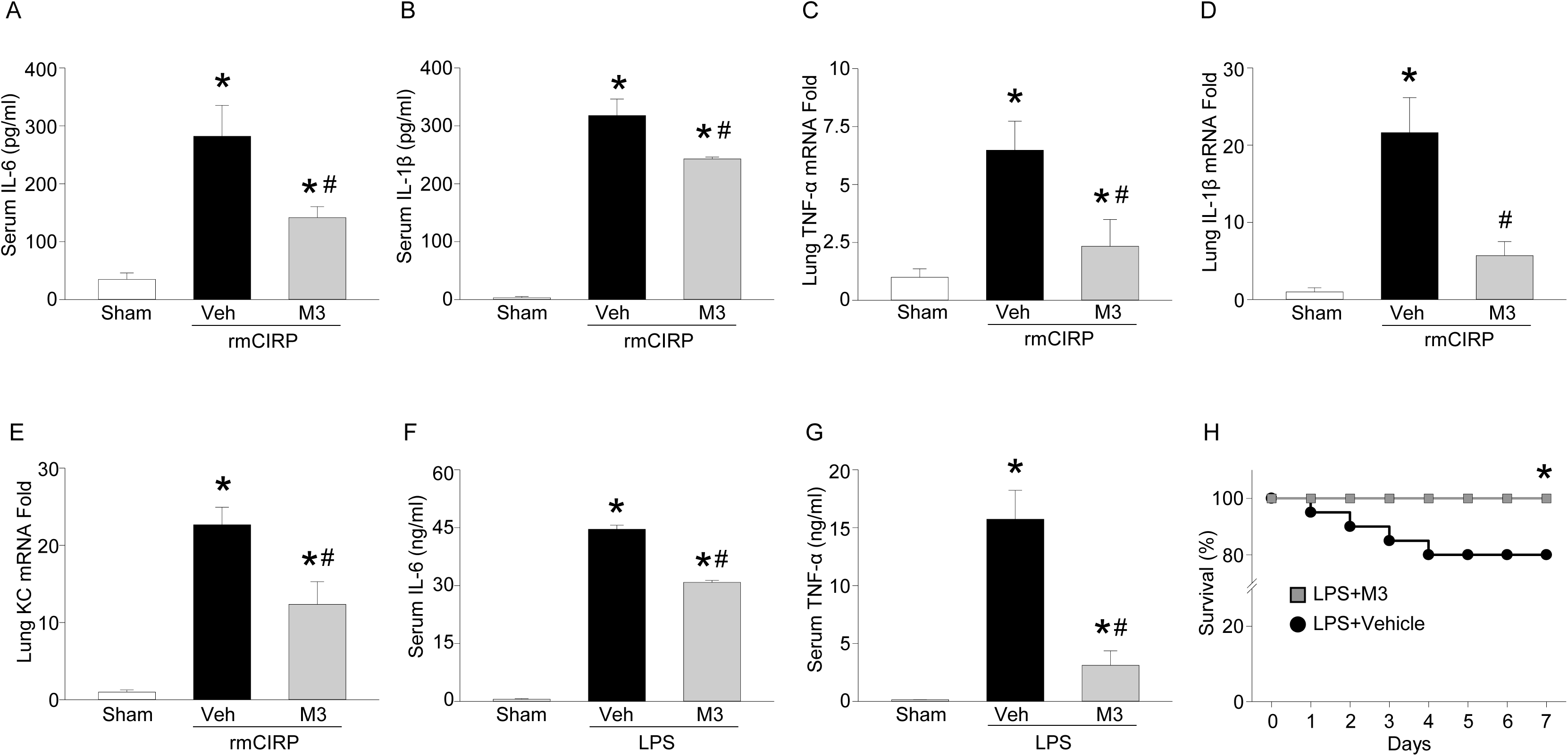
M3 inhibits eCIRP and LPS mediated inflammation. Adult C57BL/6 mice were randomly assigned to sham, vehicle (PBS), or treatment group. rmCIRP at a dose of 5 mg/kg BW or equivalent volume normal saline was administered *i.v.* via retro-orbital injection. M3 at a dose of 10 mg/kg BW or vehicle was given *i.p.* at the time of rmCIRP injection. At 5 h after rmCIRP injection, mice were euthanized, and blood and tissue were collected for analysis. Serum **(A)** IL-6 and **(B)** IL-1β were measured by ELISA. Lung mRNA levels of **(C)** TNF-α, **(D)** IL-1β, and **(E)** KC were measured by RT-PCR. Data are expressed as means ± SE. n = 4-5 mice/group. The groups were compared by one-way ANOVA and SNK method (*p<0.05 vs. sham and #p<0.05 vs. vehicle mice). Adult C57BL/6 mice were given LPS at a dose of 15 mg/kg BW or equivalent volume normal saline (sham) *i.p.* M3 at a dose of 10 mg/kg BW or vehicle (PBS) was given simultaneously. After 90 min serum was collected and ELISA was used to measure **(F)** IL-6 and **(G)** TNF-α. Data are expressed as means ± SE. n = 5 mice/group. The groups were compared by one-way ANOVA and SNK method (*p<0.05 vs. sham and #p<0.05 vs. vehicle-treated mice). C57BL/6 mice were injected *i.p.* with 15 mg/kg BW LPS and simultaneously given *i.p.* M3 at a dose of 10 mg/kg BW or equivalent volume vehicle (PBS). **(H)** Mice were monitored for survival for 7 days. n = 20 mice/group, * p<0.05 vs. LPS + vehicle (PBS), Log-rank (Mantel-Cox) test. IL, interleukin; TNF, tumor necrosis factor; ELISA, enzyme-linked immunosorbent assay; PBS phosphate buffered saline; KC, keratinocyte chemoattractant; BW, body weight; ELISA, enzyme-linked immunosorbent assay.

**Figure 4:**
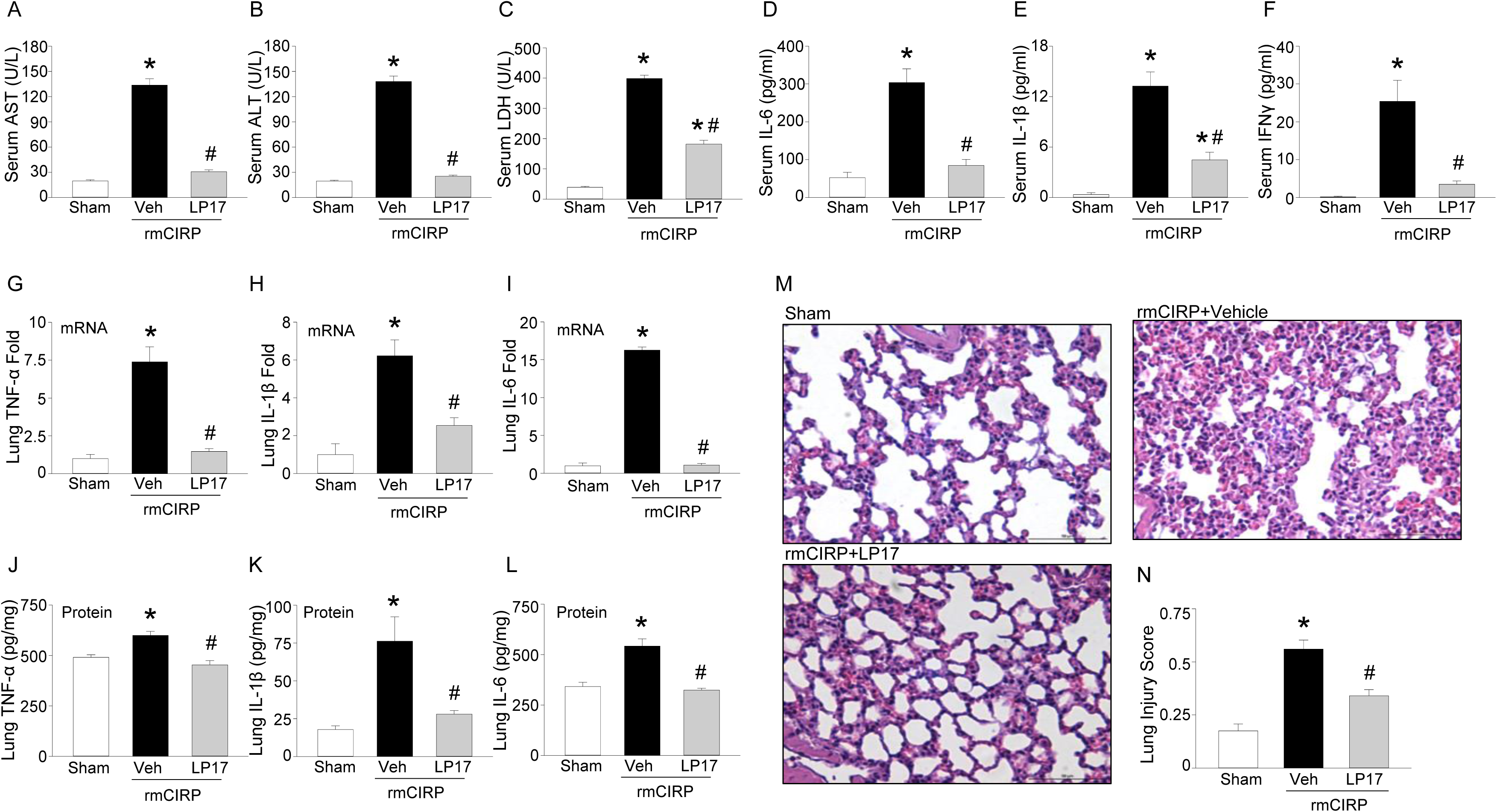
LP-17 inhibits eCIRP-induced inflammation *in vivo*. Adult C57BL/6 mice were randomly assigned to sham, vehicle (PBS), or treatment group. rmCIRP at a dose of 5 mg/kg BW or equivalent volume normal saline was administered *i.v.* via retro-orbital injection. LP17 at a dose of 5 mg/kg BW or vehicle was given *i.p.* at the time of rmCIRP injection. At 5 h after rmCIRP injection, mice were euthanized, and blood and tissue were collected for analysis. **(A)** AST, **(B)** ALT, and **(C)** LDH were determined using specific colorimetric enzymatic assays. Serum **(D)** IL-6, **(E)** IL-1β, and **(F)** IFN-γ were measured by ELISA. Lung mRNA levels of **(G)** TNF-α, **(H)** IL-1β, and **(I)** IL-6 were measured by RT-PCR. Equal amount of total lung protein (250-350 µg) were loaded into respective ELISA wells for assessment of lung protein levels of **(J)** TNF-α, **(K)** IL-1β, and **(L)** IL-6. **(M)** Representative images of H&E stained lung tissue at 200X. **(N)** Lung injury score calculated at 400x. n = 5 high powered fields/group. Data are expressed as means ± SE. n = 5-8 mice/group. The groups were compared by one-way ANOVA and SNK method (*p<0.05 vs. sham and #p<0.05 vs. vehicle mice). ALT, alanine aminotransferase; AST, aspartate amino transferase; LDH, lactate dehydrogenase; IL, interleukin; TNF, tumor necrosis factor; ELISA, enzyme-linked immunosorbent assay, H&E hematoxylin and eosin; IFN, interferon; PBS phosphate buffered saline; BW, body weight.

### M3 protects mice from polymicrobial sepsis

We next evaluated the efficacy of M3 in a more clinically relevant model of sepsis utilizing CLP in mice. CLP caused robust increases in organ injury markers AST and LDH in the serum, while the M3 treated mice showed significant decrease in their levels by 17% and 53%, respectively (Fig 5A, B). Similarly, the serum IL-6 and TNF-α were elevated by CLP, however M3 treatment significantly reduced these levels by 84% and 27%, respectively (Fig 5C, D). Expression of the pro-inflammatory cytokines IL-6, TNF-α, and chemokine KC mRNA in lung tissues were increased in CLP-induced sepsis, and were dramatically reduced with M3 treatment (Fig 5E-G). Histological images of lung tissue in CLP mice displayed significant damage, with increased levels of alveolar congestion, proteinaceous debris, interstitial and alveolar neutrophil infiltration, intra-alveolar capillary hemorrhages, and damage of epithelial architecture (Fig 5H). M3 treatment dramatically improved these histological injury parameters in septic mice (Fig 5H). These histological changes were reflected in a significant decrease in lung tissue injury score in M3-treated mice compared to vehicle mice (Fig 5I). To determine if M3 was able to improve survival in sepsis, mice were subjected to CLP and randomized to treatment with M3 or vehicle. We found that M3 treatment increased the survival rate from 45% to 80% at day 10 after CLP (Fig 5J). In summary, the released eCIRP in sepsis binds to TREM-1 and potentiates the expression of pro-inflammatory cytokines. Administration of M3, by inhibiting the eCIRP-TREM-1 interaction, exhibits excellent therapeutic potential against murine polymicrobial sepsis.

**Figure 5:**
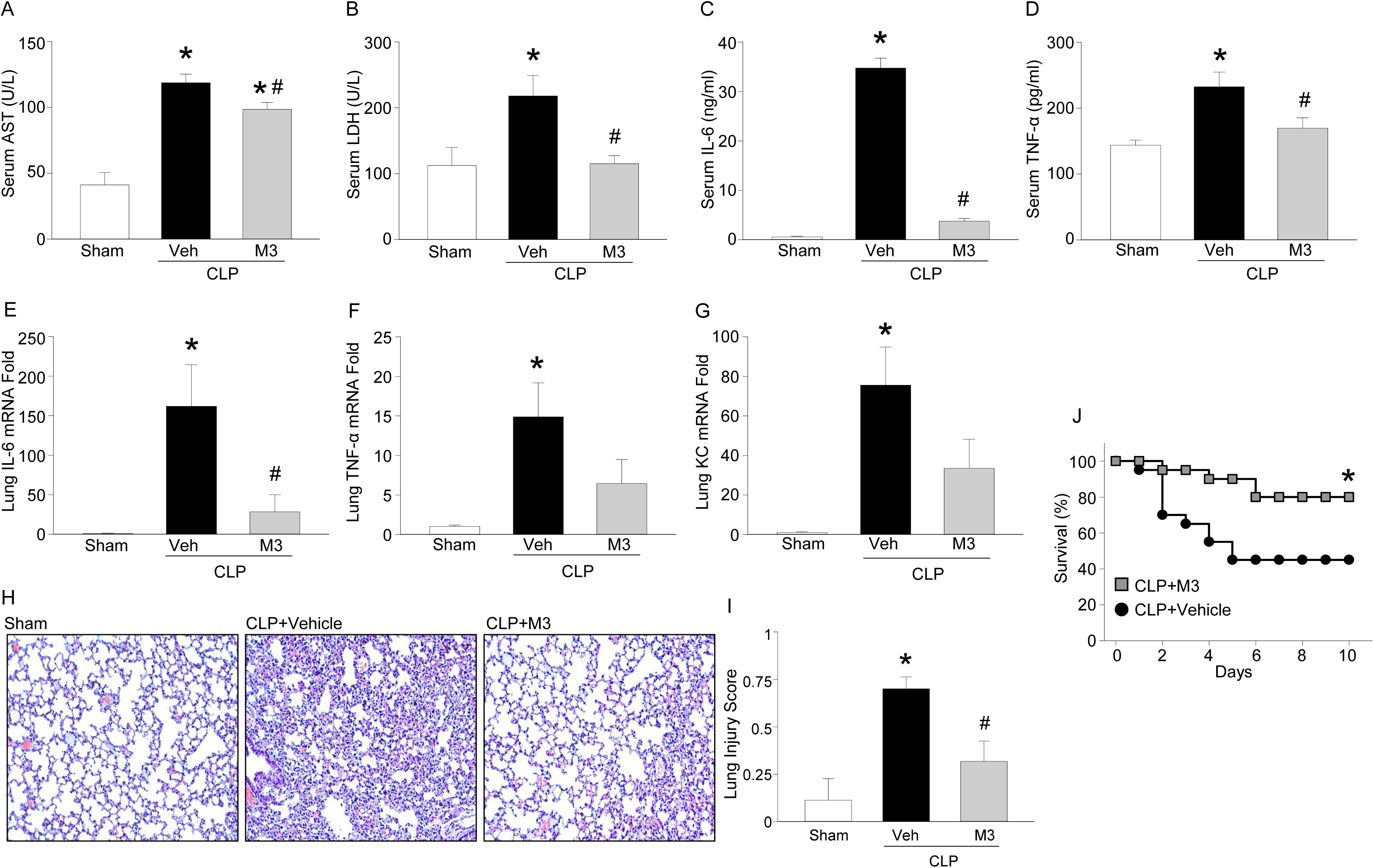
M3 protects mice from polymicrobial sepsis. Adult C57BL/6 mice were randomly assigned to sham laparotomy, CLP + vehicle (PBS), or CLP + treatment group. At the time of CLP treatment mice received an intraperitoneal instillation of 10 mg/kg body weight (BW) M3 at the time of abdominal closure. Vehicle groups received an equivalent volume of PBS. After 20 h, serum and tissue were collected for analysis. **(A)** AST and **(B)** LDH were determined using specific colorimetric enzymatic assays. Serum **(C)** IL-6 and **(D)** TNF-α were measured by ELISA. Lung mRNA levels of **(E)** IL-6, **(F)** TNF-α, and **(G)** KC were measured by RT-PCR. Data are expressed as means ± SE (n=5-8 mice/group) and compared by one-way ANOVA and SNK method (*p<0.05 vs. sham and #p<0.05 vs. vehicle-treated mice). **(H)** Representative images of H&E stained lung tissue at 200X. **(I)** Lung injury score calculated at 400x. n = 5 high powered fields/group. Data are expressed as means ± SE and compared by one-way ANOVA and SNK method (*p<0.05 vs. sham and #p<0.05 vs. vehicle mice). **(J)** Kaplan-Meier survival curve generated from treatment (M3) and vehicle CLP mice during the 10-day monitoring period is shown. n=20 mice in each group, *p<0.05 vs. vehicle, determined by the log-rank test. CLP, cecal ligation puncture; AST, aspartate amino transferase; LDH, lactate dehydrogenase; IL, interleukin; TNF, tumor necrosis factor; KC, keratinocyte chemoattractant; ELISA, enzyme-linked immunosorbent assay.

### M3 protects mice against sterile inflammation caused by intestinal I/R

As eCIRP is known to be involved in systemic and pulmonary inflammation in ischemia reperfusion injury ^6^, we evaluated the impact of M3 treatment in a murine model of intestinal I/R injury. Intestinal I/R increased serum levels of AST by 7.9-fold, ALT 3.8-fold, LDH 57.8-fold, and lactate 3.5-fold. M3 drastically improved these organ injury markers with decreases of 50%, 66%, 417%, and 47%, respectively (Fig 6A-D). Intestinal I/R increased the serum levels of IL-6 and TNF-α by 12.4 and 7.5-fold, retrospectively, while M3 treatment significantly reduced these elevations by 41% and 77%, respectively (Fig 6E, F). Lung mRNA levels of IL-6, IL-1β, and MIP-2 were significantly increased by I/R. These increases were reduced by 82%, 96%, and 89% in M3 treated I/R mice **(Supplemental Fig 5A-C)**. Protein levels of IL-6, IL-1β, and MIP-2 were increased in intestinal I/R treated mice, and M3 treatment resulted in significant reduction of 40%, 56%, and 45%, respectively (Fig 6G-I). Histological images of lung tissue showed increased levels of alveolar congestion, exudate, interstitial and alveolar cellular infiltrates, intra-alveolar capillary hemorrhages, and damage of epithelial architecture, in mice who underwent intestinal I/R compared to sham mice (Fig 6J). M3 treatment dramatically improved these histological injury parameters in intestinal I/R mice (Fig 6J). These histological changes were reflected in a significant decrease in lung tissue injury score in M3-treated compared to vehicle-treated mice by 65% (Fig 6K). M3 also doubled 24-h survival rate after I-I/R, from 40% to 80%. (Fig 6L). These results indicate that M3 is also efficacious in sterile inflammation.

**Figure 6:**
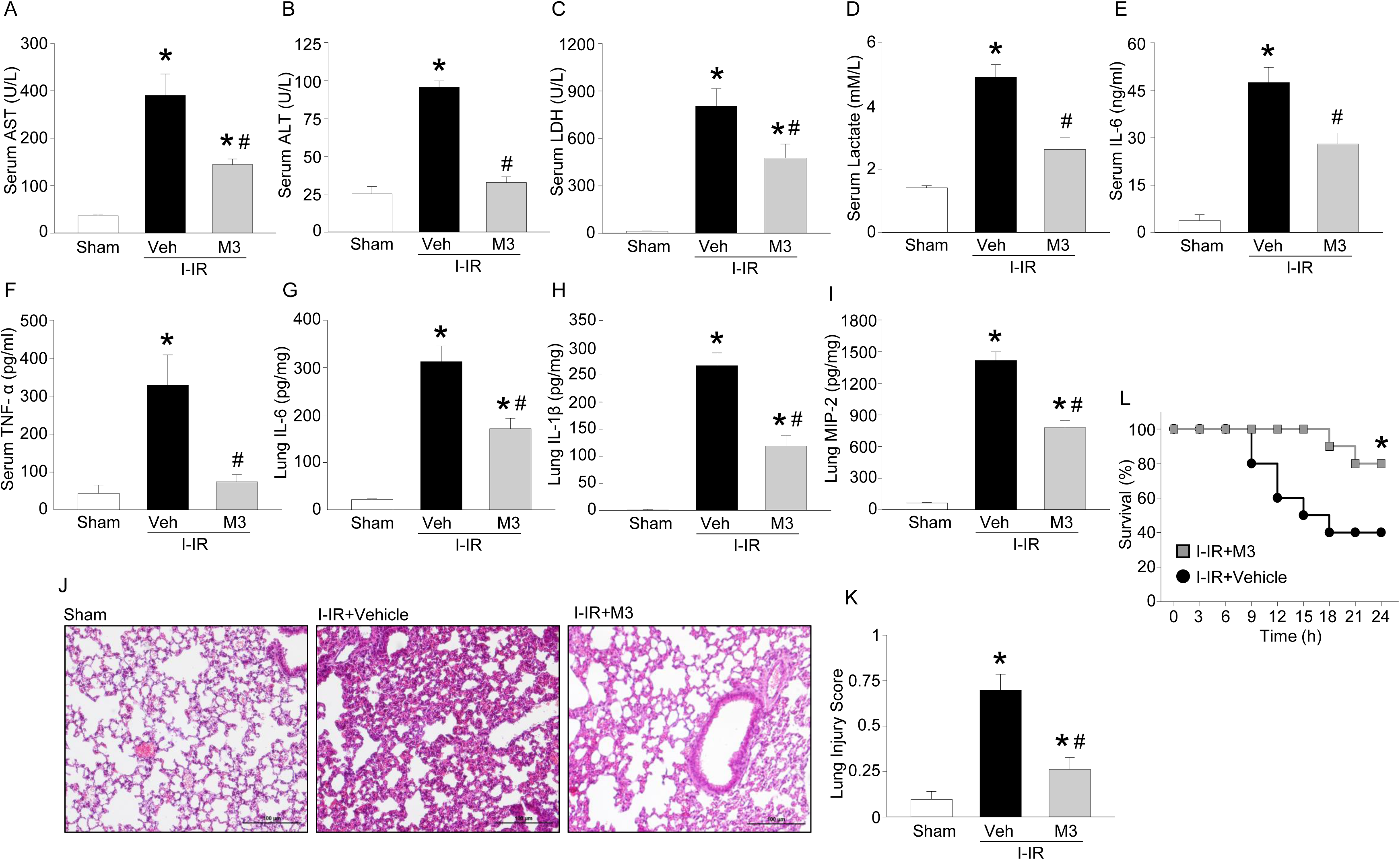
M3 protects mice against sterile inflammation caused by intestinal I/R. Mice were randomly assigned to sham laparotomy, treatment or vehicle group. I-I/R was introduced in mice via SMA occlusion for sixty minutes. Treatment mice received an intraperitoneal instillation of 10 mg/kg BW M3 at the time of reperfusion. Vehicle groups received an equivalent volume of PBS. Four hours after recovery from anesthesia, mice were sacrificed, and serum and tissue collected for analysis. **(A)** AST, **(B)** ALT, **(C)** LDH, and **(D)** lactate were determined using specific colorimetric enzymatic assays. Serum **(E)** IL-6 and **(F)** TNF-α were measured by ELISA. Equal amount of total lung protein (250-350 µg) were loaded into respective ELISA wells for assessment of lung protein levels of **(G)** IL-6, **(H)** IL-1β, and **(I)** MIP-2. **(J)** Representative images of H&E stained lung tissue at 200X. **(K)** Lung injury score calculated at 400x. n = 10 high powered fields/group. Data are expressed as means ± SE. n = 5-8 mice/group. The groups were compared by one-way ANOVA and Tukey method (*p<0.05 vs. sham and #p<0.05 vs. vehicle-treated mice). **(L)** Kaplan-Meier survival curve generated from treatment (M3) and vehicle I-I/R mice during the 24h monitoring period is shown. n=10 mice in each group, *p<0.05 vs. vehicle, determined by the log-rank test. I-I/R, intestinal ischemia-reperfusion; AST, aspartate amino transferase; ALT, alanine aminotransferase; LDH, lactate dehydrogenase; MIP-2, macrophage inflammatory protein-2; IL, interleukin; TNF, tumor necrosis factor; PBS, phosphate buffered saline; ELISA, enzyme-linked immunosorbent assay.

## Discussion

Extracellular CIRP has been recently identified as a DAMP that is released during sepsis, shock, and ischemia/reperfusion injury ^5, 6^. However, its molecular mechanism to induce inflammation still remains enigmatic. Despite identification of the TREM-1 receptor nearly two decades ago ^13^, its ligand(s) are not well elucidated. Our study fills a significant previous knowledge gap by establishing a novel link between eCIRP and TREM-1 demonstrated by several *in vitro* studies using murine and primary human macrophages and *in vivo* models clinically relevant to both sterile and infectious inflammatory diseases. We summarized the overall findings in Fig 7, which demonstrated that eCIRP released during sepsis or I/R injury binds to TREM-1, serving as a novel biologically active endogenous TREM-1 ligand. This binding leads to the activation of intracellular DAP12 and Syk and increased production of inflammatory mediators to cause hyperinflammation and tissue injury. Targeting the interaction between eCIRP and TREM-1 by a small peptide M3 derived from human eCIRP is protective against sepsis and intestinal I/R.

**Figure 7:**
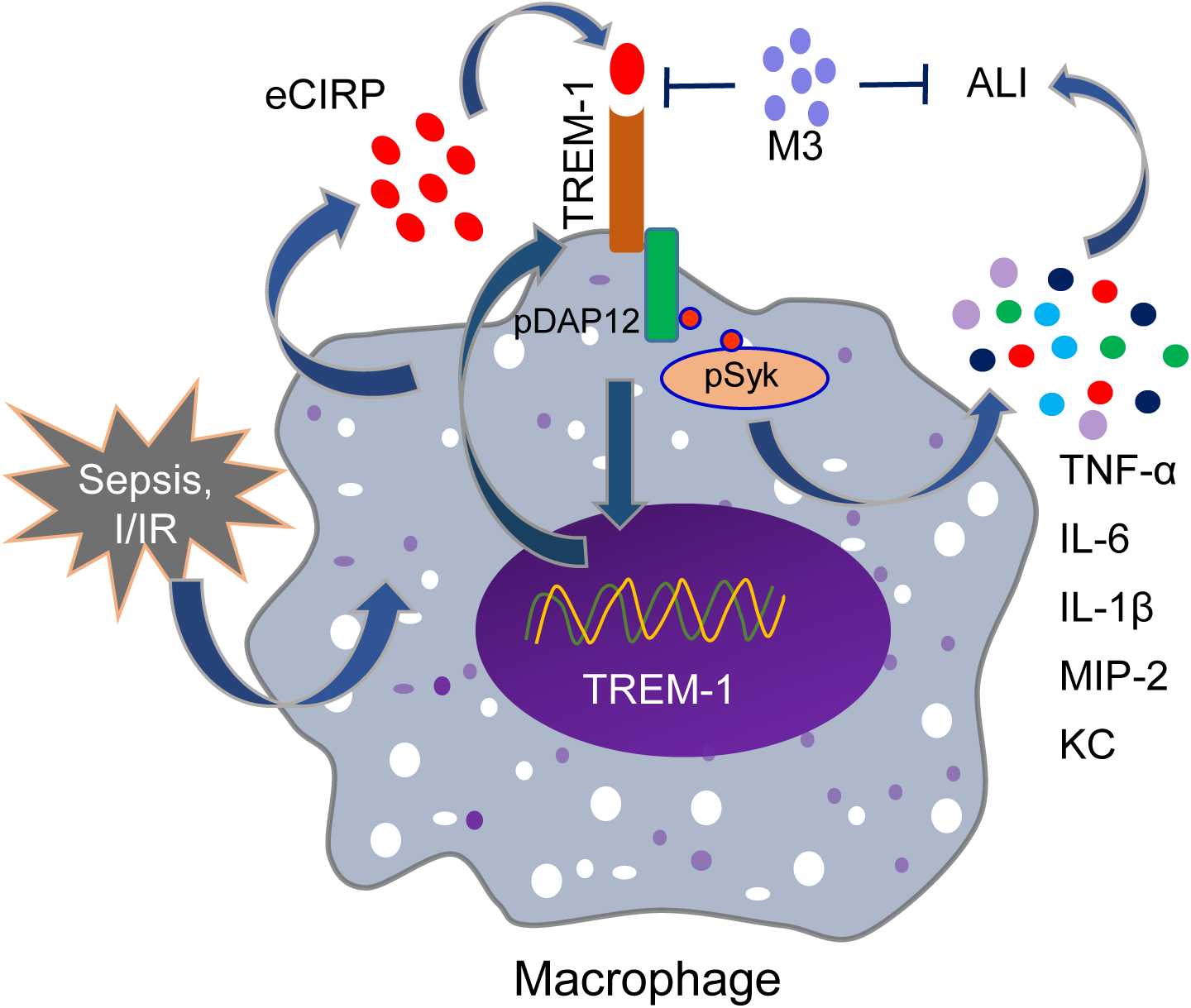
Summary of findings. Sepsis and I/R causes an increased release of eCIRP. As the endogenous ligand eCIRP recognizes TREM-1 and activates intracellular signaling molecules DAP12 and Syk, leading to increased expression of pro-inflammatory mediators that cause excessive inflammation and remote tissue injury. eCIRP increases TREM-1 expression, possibly via positive feedback induction. A small peptide M3 derived from human eCIRP abrogates eCIRP-TREM-1 interaction, thereby leading to decreased inflammation and attenuated ALI. I/R, ischemia and reperfusion; DAP12, DNAX activation protein of 12kDa; ALI, acute lung injury.

We have, for the first time, identified eCIRP as an endogenous ligand for TREM-1. In other word, TREM-1 is a new receptor for eCIRP. Previously, TLR4 was the only identified receptor for eCIRP ^5, 6^. We hypothesized that, like most DAMPs, eCIRP has several receptors for signal transduction. HMBG1, for example, is known to potentiate signals through TLR2, TLR4, receptor for advanced glycation end product (RAGE), TREM-1, and CD163 ^23, 24, 25^. Using SPR we found a strong KD of 11.7 x 10^-8^ M of binding between eCIRP and TREM-1, which is comparable to the dissociation constant of the previously identified receptor for eCIRP, TLR4 ^5^. Of the known ligands for TREM-1: PGLYRP1 ^19^, extracellular actin ^18^, and HMGB1 ^17^, SPR has been used between TREM-1 and PGLYRP1 and HMGB1. Surprisingly, the K_D_ between eCIRP and TREM-1 (11.7 × 10^-8^ M) was more than two orders lower than the KD between HMGB1 and TREM-1 (35.4 x 10^-6^ M) ^17^, indicating higher binding affinity of eCIRP to TREM-1 than between HMGB1 and TREM-1. In addition to SPR, several additional approaches were taken to confirm the physical interaction of eCIRP and TREM-1, including confocal microscopy and FRET analysis.

After demonstrating eCIRP’s binding to TREM-1, the focus was shifted to determining if eCIRP’s binding to TREM-1 resulted in fueling inflammation and causing tissue injury. This was a crucial step, as we previously demonstrated that although SPR displays binding of eCIRP to TLR2 and RAGE, there are no functional outcomes of this association ^5^. Inhibition of TREM-1 by siRNA showed some, but not complete, reduction in the production of pro-inflammatory cytokine by the macrophages treated with eCIRP. This could be due to off target effects of TREM-1 siRNA on other TREMs, although this remains speculative. TREM family contains TREM-1, TREM-2, and, in mice, TREM-3 ^26^. Although TREM-1 contains an immunoreceptor tyrosine-based activation motif (ITAM), additional TREM family members contain an immunoreceptor tyrosine-based inhibitory motif (ITIM), and thus they could play anti-inflammatory role ^27, 28^. In addition, macrophages deficient in TREM-2 are hyperresponsive to TLR ligands ^29^. We therefore used another approach to inhibit TREM-1 by LP17 which abrogated eCIRP-induced inflammation and ALI, confirming the notion that the eCIRP-TREM-1 interaction is functionally dynamic. LP17 is a 17 amino acid peptide whose sequence is taken from TREM-1; it is thought to function as a decoy receptor ^21^. We attempted to determine the interaction between LP17 and eCIRP using SPR and demonstrated a KD of 48 x 10^-6^ M between these two molecules (data not shown). This could be artificially elevated due to conditions present *in vivo* that we cannot reproduce artificially, such as involvement of other biological co-factors, temperature, and molecular stability. The TREM-1 antagonists function primarily as decoy receptors and to prevent receptor dimerization ^21, 30^. Additionally, Sigalov *et al* has developed a ligand-independent inhibitor of TREM-1 ^31, 32^. Although use of these peptides in animal models of inflammatory diseases has resulted in decreased inflammation and improved survival, it is not clear which ligand of TREM-1 these decoy peptides neutralize, rendering their use somewhat less specific. Here, we have developed a ligand dependent TREM-1 inhibitor, a novel short peptide M3. As demonstrated with FRET analysis and functionally *in vitro* and *in vivo*, M3 is able to prevent eCIRP’s interaction with TREM-1 thereby impeding eCIRP’s pro-inflammatory role.

Both the specificity to the eCIRP and TREM-1 interaction and the small size of M3 may prove advantageous in future therapeutic applications of M3. M3 has a molecular weight of 795.9 Daltons. Most peptides in clinical use are injected subcutaneously. Peptides under 1 kDa can be absorbed directly into the systemic circulation via blood capillaries, whereas larger peptides are absorbed initially through the lymphatic system or in a combination of both blood and lymphatic absorption ^33, 34^. Additionally, because patient compliance is increased with non-invasive administration, small peptides have an advantage over larger peptides. In the current study, we demonstrated M3 had an excellent inhibitory role in mitigating inflammation not only in macrophage cells, but also in bacterial sepsis and sterile inflammation. Prior to usage of M3 in large animal models or human clinical trials, further studies on the stability, half-life, and safety would be required. However, at the doses used here, M3 did not demonstrate immunogenicity or tissue injury as demonstrated by cytokine levels, organ injury markers, or histologic analysis.

In addition to eCIRP’s interaction with TREM-1, we also found that the expression of TREM-1 in macrophages was increased following treatment with eCIRP. Several studies have established the connection between TLR4 and TREM-1 in immune cells ^14, 35, 36^. The cellular consequences of TREM-1 activation following treatment with anti-TREM-1 Ab were examined in terms of global gene expression and compared with cells receiving LPS and also cells receiving a combined treatment of anti-TREM-1 plus LPS ^14^. The results indicated that besides having the crosstalk with TLR4, TREM-1 signaling independently promoted the expression of some unique genes ^14^, implying the direct impact of TREM-1 pathway in inflammation. As previously discussed, like other DAMPs, eCIRP has been shown to recognize TLR4 ^5^. TLR4 activation leads to upregulation of TREM-1 expression which is MyD88-dependent and involves transcription factors NF-κB, PU.1 and AP1 ^37^. Additionally, simultaneous activation of TREM-1 and TLR4 leads to synergistic production of pro-inflammatory mediators through common signaling pathway activation including PI3K, ERK1/2, IRAK1 and NF-κB activation ^13, 14, 15, 16^. Thus, the upregulation of TREM-1 expression in macrophages after treatment with eCIRP could be promoted through both TLR4 and TREM-1-dependent pathways. The connection between TLR4 and TREM-1 has been shown in neutrophils: following LPS stimulation of neutrophils, TREM-1 was found to be co-localized with TLR4 ^36^. Genetic deletion of TREM-1 down-regulates the expression of several genes implicated in the TLR4 pathway including MyD88, CD14 and IκBα ^35^. Our data demonstrating the eCIRP-induced expression of pro-inflammatory cytokines by the macrophages also supports the synergistic effects of TREM-1 and TLR4. However, to demonstrate the independent pro-inflammatory role of the TREM-1 in eCIRP-mediated inflammation, we activated TREM-1 using agonist antibody crosslinking ^13, 14^. TREM-1 activation resulted in increased levels of TNF-α; however, TNF-α production was diminished in cells treated with M3. M3 interfered with the crosslinking effect demonstrating a TLR4 independent interaction between eCIRP and TREM-1.

In summary, we have discovered that eCIRP is a new ligand of TREM-1 in macrophages. We have developed a small peptide, M3, which successfully reduces systemic inflammation and improves survival in murine models of infectious and sterile inflammation. M3 was able to successfully inhibit inflammation in a human cell line. Future work should focus on additional immune cell types and other models of inflammatory diseases. Identification of eCIRP as a new biologically active endogenous TREM-1 ligand has allowed additional insight into the pathobiology of eCIRP-mediated inflammation in sepsis and intestinal I/R.

## Methods

### Experimental animals

C57BL/6 male mice were purchased from Charles River Laboratories (Wilmington, MA). Age (8-12 weeks) matched healthy mice were used in all experiments. Animals were randomly assigned to sham, vehicle, or treatment group. The number of animals used in each group was based on our previous publication ^38^. All experiments were performed in accordance with the guidelines for the use of experimental animals by the NIH and were approved by the Institutional Animal Care and Use Committee.

### Peptides

LP17 (LQVTDSGLYRCVIYHPP), LP17-scramble control (TDSRCVIGLYHPPLQVY), M3 (RGFFRGG) and M3-scramble controls: M3-Sc1 (FGRGFRG) and M3-Sc2 (GFFGRGR) were synthesized by GenScript (Piscataway, NJ).

### Animal model of polymicrobial sepsis

Mice were anesthetized with inhaled isoflurane and placed in supine position. Cecal ligation and puncture (CLP) was preformed through a midline laparotomy ^39, 40^. Briefly, the abdomen was shaved and disinfected. A 2-cm incision was created, the cecum was exposed, and ligated with a 4-0 silk suture 1 cm proximal from the distal cecal extremity. For 20 h experiments the cecum was punctured twice with a 22-guage needle. For the 10-day CLP experiments the cecum was only punctured once with the same sized needle. A small amount of cecal content was extruded and the ligated cecum was returned to the peritoneal cavity. The wound was closed in layers. Mice were allocated to treatment or vehicle group. Treatment mice received an intraperitoneal injection of 10 mg/kg body weight (BW) M3 at the time of abdominal closure. Vehicle groups received an equivalent volume of phosphate buffered saline (PBS). Sham animals underwent a laparotomy without cecal ligation or puncture. After closure, the mice received a subcutaneous injection of 1 ml of normal saline to avoid surgery-induced dehydration. Animals in the 20 h experiment were not given antibiotics in order for mice to develop severe sepsis and early mortality. Animals in the 10-day survival experiment were given 500 μl of the antibiotic Imipenem (0.5 μg/kg, Merck, Kenilworth, NJ) and 500 μl of normal saline subcutaneously at the time of laparotomy. For the 10-day experiment, mice were evaluated twice daily for their survival status.

### Animal model of endotoxemia

Mice were injected intraperitoneally (*i.p.*) with 15 mg/kg BW of LPS (Escherichia coli serotype O55:B5; Sigma-Aldrich, St Louis, MO). Mice were allocated to treatment or vehicle group. Treatment mice received an *i.p.* injection of 10 mg/kg BW M3 at the time of LPS injection. Vehicle group received an equivalent volume of PBS. After 90 min, mice were sacrificed, and serum was collected for analysis. To study survival rates, mice were treated with 15 mg/kg BW LPS (Escherichia coli serotype O55:B5; Sigma-Aldrich) with or without 10 mg/kg BW M3 and were evaluated twice daily for 7 days.

### Animal model of intestinal ischemia/reperfusion

Mice were anesthetized with inhaled isoflurane and the abdomen was shaved and disinfected. Similar to previous descriptions ^41^, an upper midline laparotomy was performed, and the superior mesenteric artery was isolated. The superior mesenteric artery was occluded with a vascular clip resulting in 60 min of ischemia time. After 60 min, the vascular clip was removed to allow reperfusion. Mice were randomly allocated to treatment or vehicle group. Treatment mice received an intraperitoneal instillation of 10 mg/kg BW M3 at the time of reperfusion. Vehicle groups received an equivalent volume of PBS. Sham animals underwent a laparotomy with the same 1 h of anesthesia time without arterial occlusion. The abdomen was closed in layers and mice were resuscitated with 1 ml of normal saline injected subcutaneously. At 4 h after reperfusion mice were euthanized and serum and tissue collected for analysis. For the survival study, mice were evaluated every 4 h for 24 h.

### *In vivo* administration of rmCIRP, LP17 and M3

rmCIRP was produced in our lab as previously described ^5^. rmCIRP at a dose of 5 mg/kg BW or normal saline was administered intravenously (*i.v.*) via retro-orbital injection using an 29G × 1/2″ U-100 insulin syringe (Terumo Medical Corporation, Elkton, MD). LP17 at a dose of 5 mg/kg BW, or M3 (10 mg/kg BW) or vehicle (PBS) was given *i.p.* at the time of rmCIRP injection. At 5 h after rmCIRP injection, mice were anesthetized, and blood and tissue were collected for analysis. A section of lung tissue was preserved in 10% formalin for histopathological analysis and rest was frozen in liquid nitrogen and stored at −80 °C for qPCR analysis.

### Cell culture and isolation of peritoneal macrophages

Mouse macrophage RAW264.7 cells were obtained from ATCC (Manassas, VA). Macrophages from healthy human donors were obtained from Hemacare (Cat. No. PBMACC-MON-2, Northridge, CA). Murine peritoneal macrophages were isolated from adult male mice^38^. Briefly, 1 ml of 4% thioglycolate was injected intraperitoneally. Four days later, mice were euthanized using CO2 asphyxiation. Peritoneal fluid and cells were isolated using peritoneal lavage with PBS. Total peritoneal cells were isolated by centrifugation at 200g for 10 min and subsequently cultured in Dulbecco’s Modified Eagle’s Medium (DMEM) (Thermo Fischer-Scientific). After 4 h nonadherent cells were removed and adherent cells, primarily macrophages, were cultured overnight prior to use. All cultured media was supplemented with 10% heat-inactivated fetal bovine serum (FBS, MP Biomedicals, Solon, OH), 1% penicillin-streptomycin and 2 mM glutamine. Cells were maintained in a humidified incubator with 5% CO_2_ at 37 °C.

### Determination of organ injury markers

Serum levels of lactate dehydrogenase (LDH), aspartate aminotransferase (AST), alanine aminotransferase (ALT), and lactate were determined using specific colorimetric enzymatic assays (Pointe Scientific, Canton, MI) according to the manufacturer’s instructions.

### Enzyme-linked immunosorbent assay (ELISA)

Supernatant or serum was analyzed by ELISA kits specific for interleukin (IL)-6, tumor necrosis factor-α (TNF-α), interferon-γ (IFN-γ), (BD Biosciences, San Jose, CA), IL-1β (Invitrogen, Carlsbad, CA), and macrophage inflammatory protein-2 (MIP-2) (R&D Systems, Minneapolis, MN) according to the manufacturer’s instructions. The lung tissue was crushed in liquid nitrogen, and equal weights of powdered tissues (∼50 mg) were dissolved in 500 µl of lysis buffer (10 mM Hepes, pH 7.4, 5 mM MgCl2, 1 mM DTT, 1% Triton X-100, and 2 mM each of EDTA and EGTA), and subjected to sonication on ice. Protein concentration was determined by the BioRad protein assay reagent (Hercules, CA). Equal amounts of proteins (250-500 µg) were loaded into respective ELISA wells for the assessment of TNF-α, IL-6, MIP-2, and IL-1β.

### Lung pathohistology

Lung tissues were fixed in 10% formalin prior to being embedded in paraffin. Tissues were cut into 5 μm sections and stained with hematoxylin and eosin (H&E). Slides were evaluated under light microscopy to evaluate the degree of lung injury. Scoring was done using a system created by the American Thoracic Society ^42^. Scores ranged from zero to one and were based on the presence of proteinaceous debris in the airspaces, the degree of septal thickening, and neutrophil infiltration in the alveolar and interstitial spaces. The average score per field was calculated at 400× magnification.

### Assessment of TREM-1 expression in macrophages by flow cytometry

To detect TREM-1 expression on the surface of macrophages, a total of 1 × 10^6^ RAW264.7 or primary peritoneal macrophages were washed with FACS buffer containing PBS with 2% FBS and stained with APC anti-mouse TREM-1 Ab (clone: 174021, R&D systems). Unstained cells were used as a negative control to establish the flow cytometer voltage setting. Acquisition was performed on 10,000 events using a BD LSR Fortessa flow cytometer (BD Biosciences, San Jose, CA) and data were analyzed with FlowJo software (Tree Star, Ashland, OR).

### Real-time quantitative reverse transcription polymerase chain reaction (qRT-PCR)

Total RNA was extracted from tissue using Trizol reagent (Invitrogen, Carlsbad, CA). cDNA was synthesized using MLV reverse transcriptase (Applied Biosystems, Foster City, CA). PCR reactions were carried out in 20 μl of a final volume of 0.08 μM of each forward and reverse primer (**Supplemental Table 1**), cDNA, water, and SYBR Green PCR master mix (Applied Biosystems). Amplification and analysis was conducted in a Step One Plus real-time PCR machine (Applied Biosystems). Mouse β-actin mRNA was used as an internal control for amplification and relative gene expression levels were calculated using the ΔΔCT method. Relative expression of mRNA was expressed as fold change in comparison with sham tissues.

### Immunoprecipitation and Western blotting

For immunoprecipitation assay of DAP12 and phosphotyrosine, RAW264.7 cells (1 × 10^6^) were stimulated with rmCIRP (1 µg/ml). At different time points, cells were harvested and proteins were extracted using extraction buffer containing 25 mM Tris, 0.15 M NaCl, 1 mM EDTA, 1% NP-40, 5% glycerol, 2 mM Na_3_VO_4_, and protease inhibitor cocktail (Roche Diagnostics, Indianapolis, IN), pH 7.4. Equal amounts of protein were subjected to do immunoprecipitation using anti-DAP12 Ab (Cell Signaling Technology, Danvers, MA) and protein A/G plus agarose beads (Pierce Classic IP Kit, Thermo Fischer Scientific). Following immunoprecipitation, samples were eluted and run on 4-12% gradient polyaccryl amide gel for electrophoresis. Gels were transferred into nitrocellulose membranes, blocked with 3% BSA and finally reacted with primary antibodies for phosphotyrosine (Cat No.: 05-321; clone: 4G10; EMD Millipore, Burlington, MA), DAP12 (Cell Signaling Technology). For the analysis of pSyk, Syk, and β-actin, equal amounts of protein were subjected to perform Western blotting using 4-12% gradient polyaccryl amide gels. The blots were reacted with pSyk (Abcam; Cambridge, MA), Syk (Cell Signaling Technology), and β-actin (Sigma-Aldrich, St Louis, MO) primary Abs, followed by reaction with fluorescent-labeled secondary Abs (Li-Cor Biosciences, Lincoln, NE) and detection by Odyssey FC Dual-Mode Imaging system (Li-Cor Biosciences). The intensities of the bands we measured by using Image Studio Ver 5.2 (Li-Cor Biosciences).

### Stimulation of cytokine production by TREM-1 engagement

To activate RAW264.7 cells through TREM-1, 96-well flat bottom plates were pre-coated with 20 μg/ml of an agonist anti-TREM-1 mAb (clone 174031; R&D Systems) overnight at 37°C, similar to previously described protocols ^13, 15^. The wells were washed with sterile PBS and 5 × 10^4^ cells/well were plated. Cells treated with M3 or scramble M3-Sc1 were premixed with the peptide for 30 min before adding to the well. TNF-α production was measured in the culture supernatants after an additional 24 h of incubation.

### Surface plasmon resonance

To examine the direct interaction between eCIRP and TREM-1, surface plasmon resonance (SPR), using Biacore technology (3000 instrument, GE Healthcare), was performed between rmCIRP and rmTREM-1. Anti-his Ab was used to capture rmCIRP-his. TREM-1 (recombinant mouse TREM-1-Fc Chimera, R&D Systems, Cat No.: 1187-TR-025) was injected as an analyte in concentrations of 0 to 500 nM. Binding reactions were performed in 1×HBS-EP (10 mM Hepes, 150 mM NaCl, 3 mM EDTA, 0.05% P20, pH7.4) The CM5 dextran chip (flow cell-2) was first activated by injection with mix of 35 μl of 0.1 M N-ethyl-N′-[3-diethylaminopropyl]-carbodiimide and 0.1 M N-hydroxysuccinimide. An aliquot of 200 μl of 5 μg ml−1 of the ligand diluted in 10 mM sodium acetate (pH 4.5) was injected into flow cell-2 of the CM5 chip for immobilization. Next, 35 μl of 1 M ethanolamine (pH 8.2) was injected to block the remaining active sites. The flow cell-1 was treated with 35 μl of 0.1 M N-ethyl-N′-[3-diethylaminopropyl]-carbodiimide and 0.1 M N-hydroxysuccinimide and blocked with 1 M ethanolamine (pH 8.2) without the ligand and used as a control to evaluate nonspecific binding. The binding analyses were performed at a flow rate of 30 μl min−1 at 25 °C. To evaluate the binding, the analyte (0.5 μM for the yes-or-no binding analysis, and ranging from 31.25 nM to 0.5 μM for the kinetics analysis) was injected into flow cell-1 and flow cell-2, and the association of analyte and ligand was recorded by SPR. The signal from the blank channel (flow cell-1) was subtracted from the channel coated with the ligand (flow cell-2). Data were analyzed by the BIAcore 3000 Evaluation Software. For all samples, a blank injection with buffer alone was subtracted from the resulting reaction surface data. Data were globally fitted to the Langmuir model for 1:1 binding.

### Immunofluorescence assay

RAW264.7 cells were treated with rmCIRP (5 µg/ml) at 4°C for 10 min and then fixed in a nonpermeabilizing fashion using 4% paraformaldehyde. Immunofluorescence staining was performed using primary Abs against TREM1, CD11b, and CIRP and fluorescently tagged secondary antibodies. The fixed cells were washed in PBS and blocked with normal 5% normal horse serum for 1 h. Primary antibodies were diluted in 1% horse serum and incubated with cells for 2 h at room temperature or overnight at 4°C. After washing with PBS, cells were incubated in the dark with diluted secondary Abs for 1 h. After an additional washing step, slides were mounted immediately on Vectashield mounting medium with DAPI (Vector Laboratories, Burlingame, CA). Antibodies used were as follows: rabbit anti-mouse CIRP Ab (Catalog 10209-2-AP; ProteinTech, Rosemont, IL), goat anti-mouse TREM-1 Ab (Catalog AF1187; R&D Systems), goat anti-CD11b Ab (Catalog MBS420973MyBioSource, San Diego, CA), Cy5-conjugated AffiniPure donkey anti-goat IgG (Code 705-175-147; Jackson ImmunoResearch Laboratories, West Grove, PA) and Cy3-conjugated AffiniPure F(ab’)2 fragment donkey anti-rabbit IgG (Code 711-166-152; Jackson ImmunoResearch Laboratories). Confocal microscopy images were obtained at using a Zeiss LSM880 confocal microscope equipped with a 63× objective (Zeiss, Oberkochen, Germ). Images were analyzed and quantified by using the ZenBlue software (Zeiss, Oberkochen, Germany).

### FRET analyses

FRET analysis was performed as described previously ^43^. RAW264.7 cells were treated with rmCIRP at 4°C for 10 min with or without M3 or scramble peptide, fixed in a nonpermeabilized fashion, and stained using the antibodies and protocol listed under Immunofluorscence methods. Cell associated fluorescence was measured on Biotek Synergy Neo2 (Biotek, VT) at 566 nm upon excitation at 488 nm (*E*1), at 681 nm after excitation at 630 nm (*E*2), and at 681 nm after excitation at 488 nm (*E*3). The transfer of fluorescence was calculated as FRET units. FRET unit= [*E*3_both_ −*E*3_none_] − [(*E*3_Cy5_ −*E*3_none_) × (*E*2_both_/*E*2_Cy5_)] − [(*E*3_Cy3_ − *E*3_none_) × (*E*1_both_/*E*1_Cy3_)].

### Statistical analysis

Data represented in the figures are expressed as mean ± SE. ANOVA was used for one-way comparison among multiple groups and the significance was determined by the Student– Newman–Keuls (SNK) test or the Tukey method, as appropriate. The paired Student *t* test was applied for two-group comparisons. Significance was considered for *p* ≤ 0.05 between study groups. Data analyses were carried out using GraphPad Prism graphing and statistical software (GraphPad Software, San Diego, CA).

## Author contributions

DNL, MA designed the experiments. DNL performed animal and *in vitro* experiments. AM, SG performed *in vitro* mechanistic studies. MO performed animal survival studies. DNL, MA, JMP analyzed the data. DNL, MA, PW prepared the figures and wrote the manuscript. PW revised and edited the manuscript. PW conceived the idea and supervised the project.

## Acknowledgements

We thank Michael Bloch and Ying Zhang of University of Maryland and Archna Sharma of Center for Immunology and Inflammation (CII), Feinstein Institute for Medical Research (FIMR) for BIAcore assays, Amanda Chen of microscopy core, FIMR for microscopic study, Betsy Barnes, Center for Autoimmune, Musculoskeletal, and Hematopoietic Diseases, FIMR for the FRET assay help, and Max Brenner of CII, FIMR for critical review.

## Competing financial and/or non-financial interests

One of the authors (PW) is an inventor of patent applications covering the fundamental concept of targeting CIRP for the treatment of inflammatory diseases, licensed by TheraSource LLC. PW is a co-founder of TheraSource LLC. Other authors declared that they have no competing interests.

