## Supplemental Figures for "Extracellular CIRP as a Novel Endogenous TREM-1 Ligand to Fuel Inflammation"

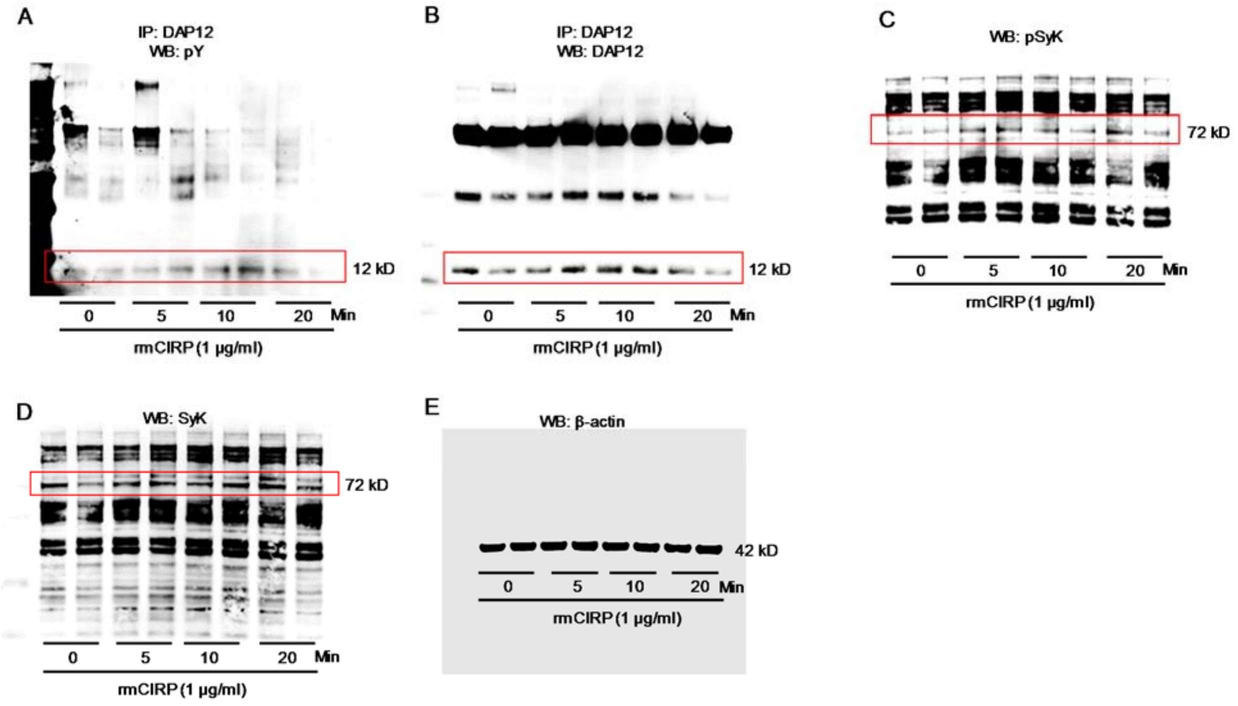


**Supplemental Figure 1: eCIRP induces TREM-1 downstream molecules**

A total of 1 × $10^6$/ml RAW264.7 cells were stimulated with rmCIRP (1 μg/ml) for various times. Proteins were immunoprecipitated by using anti-DAP12 Ab, followed by Western blotting using pTyr (4G10) and DAP12 Ab. Extracted proteins obtained from rmCIRP (1 μg/ml for various times) stimulated RAW264.7 cells (1 × $10^6$/ml) were subjected to Western blotting using pSyk, Syk, and β–actin Abs. **(A-E)** Full blots of representative Western blots for pTyr (4G10), DAP12, pSyk, Syk, and β–actin are shown. DAP12, DNAX activation protein of 12kDa; rmCIRP, recombinant mouse CIRP.

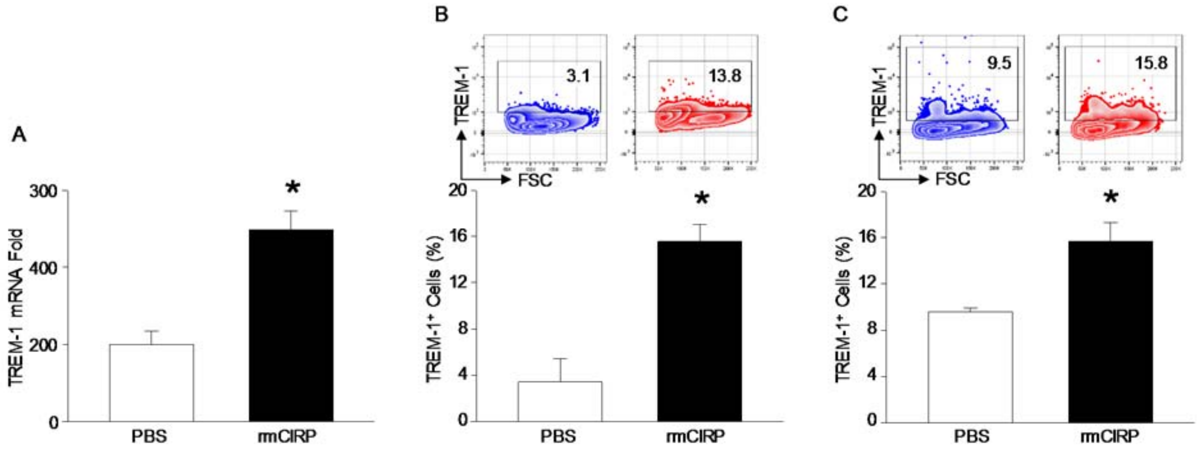


**Supplemental Figure 2: Extracellular CIRP induces TREM-1 expression in macrophages**

***(A)*** RAW264.7 cells were stimulated with PBS or rmCIRP (1 μg/ml). After 6 h, RT-PCR was done for TREM-1. Data are expressed as means ± SE, two independent experiment, n = 5/group, unpaired T-test, *p <0.05 vs PBS. **(B)** To detect TREM-1 expression on the surface of macrophages, a total of 1 × $10^6$ RAW264.7 or **(C)** primary peritoneal macrophages were stained with APC anti-mouse TREM-1 Ab. Acquisition was performed on 10,000 events using a BD LSR Fortessa flow cytometer and data were analyzed with FlowJo software. Data are expressed as means ± SE, n = 3/group, unpaired T-test, *p <0.05 vs PBS. PBS, phosphate buffered saline; APC, allophycocyanin.

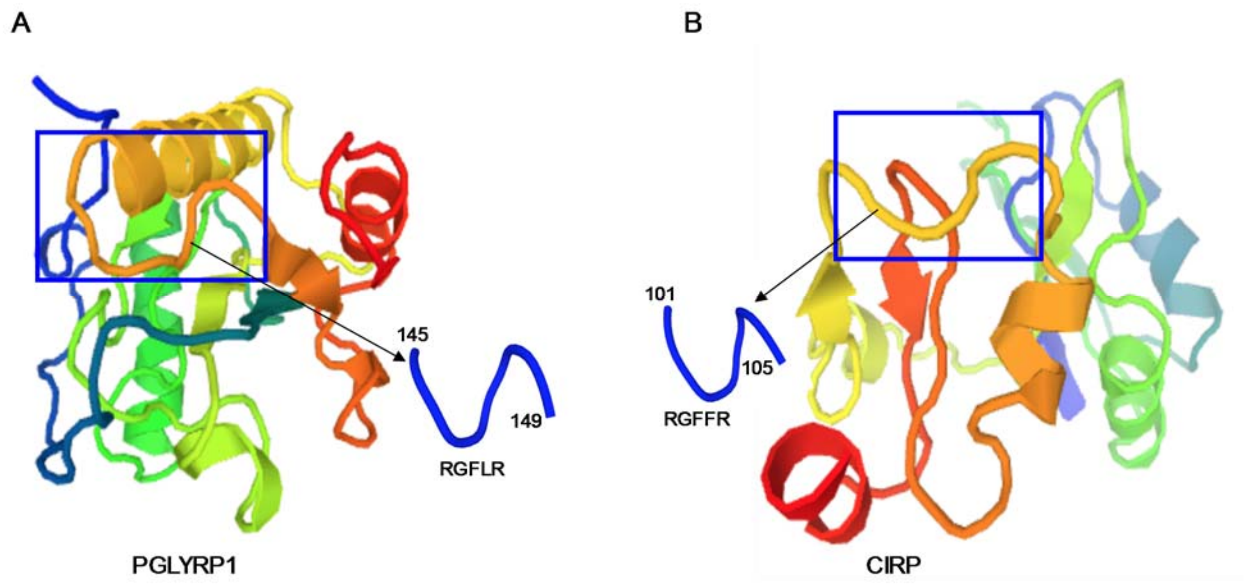


**Supplemental Figure 3: Development of the M3 peptide**

Three-dimensional structures generated using the Protein Model Portal of **(A)** murine PGLYRP1 and **(B)** murine CIRP. Murine and human CIRP share 96% amino acid sequence homology. Boxed area highlights an area of structural similarity. PGLYRP1, peptidoglycan recognition protein 1.

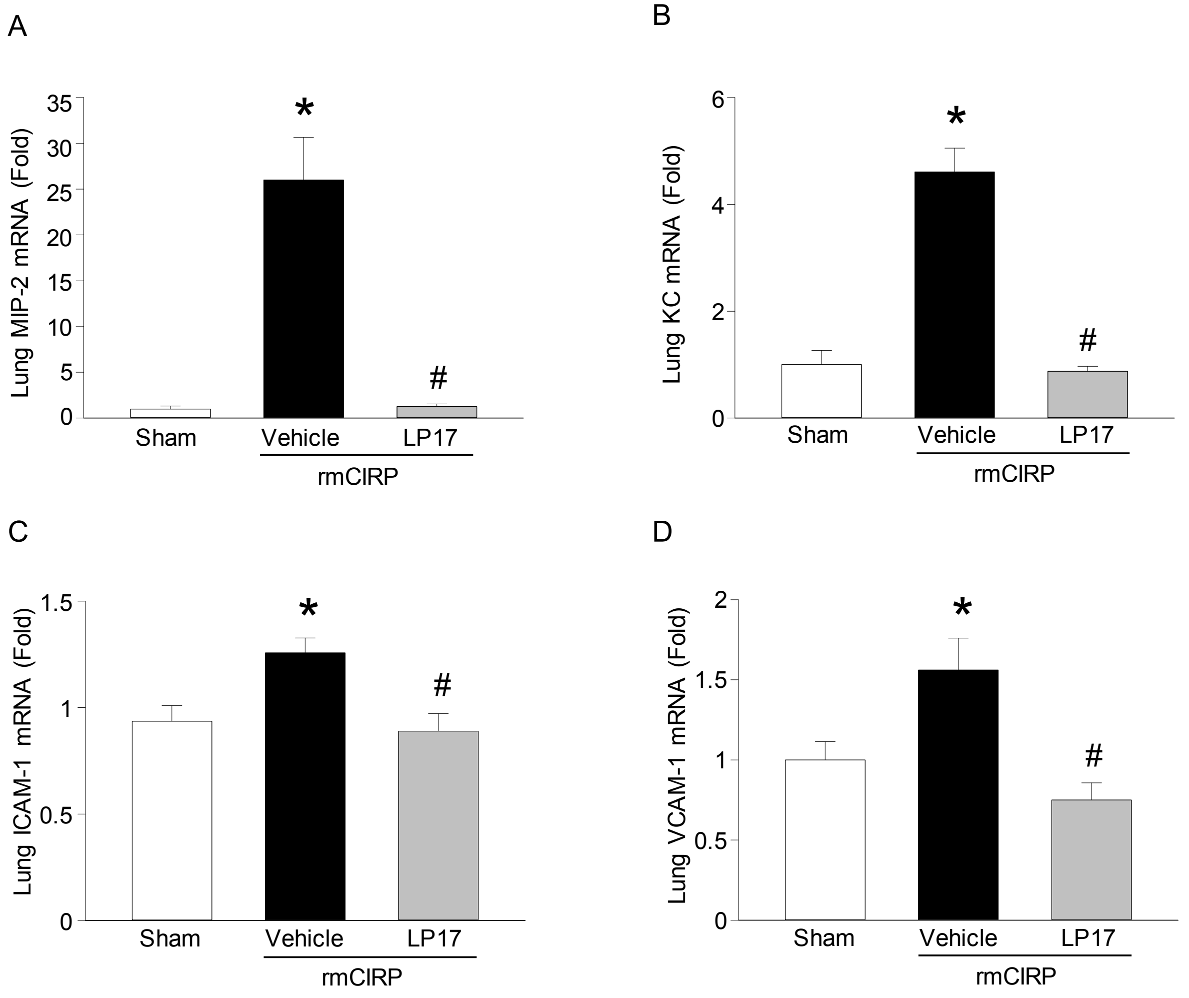


**Supplemental Figure 4: LP17 treatment inhibits lung chemokine and adhesion molecules following rmCIRP injection in mice**

Adult C57BL/6 mice were randomly assigned to sham, vehicle, or treatment group. rmCIRP at a dose of 5 mg/kg BW or equivalent volume normal saline was administered *i.v.* via retro-orbital injection. LP17 at a dose of 5 mg/kg BW or vehicle (PBS) was given *i.p.* at the time of rmCIRP injection. At 5 h after rmCIRP injection, mice were euthanized, and tissue was collected for analysis. Lung mRNA levels of **(A)** MIP-2, **(B)** KC, **(C)** ICAM-1, and **(D)** VCAM-1 were measured by RT-PCR. Data are expressed as means ± SE. n = 5-8 mice/group. The groups were compared by one-way ANOVA and SNK method (*p<0.05 vs. sham and #p<0.05 vs. vehicle-treated mice). MIP-2, macrophage inflammatory protein-2;

KC, keratinocyte chemoattractant; ICAM, intracellular adhesion molecule; VCAM, vascular cell adhesion protein.

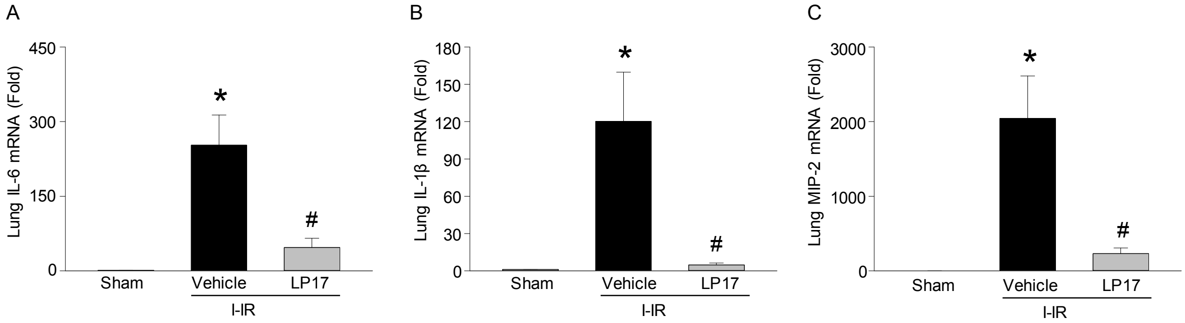


**Supplemental Figure 5: M3 inhibits cytokine and chemokine expression in the lung tissues during intestinal ischemia-reperfusion injury**

Intestinal I/R was introduced in mice via SMA occlusion for sixty minutes. Treatment mice received an intraperitoneal injection of 10 mg/kg BW M3 at the time of reperfusion. Vehicle groups received an equivalent volume of PBS. Four hours after recovery from anesthesia, mice were sacrificed, and tissue was collected for analysis. Lung mRNA levels of **(A)** IL-6, **(B)** IL-1β, and **(C)** MIP-2 were measured by RT-PCR. Data are expressed as means ± SE. n = 5-8 mice/group. The groups were compared by one-way ANOVA and Tukey method (*p<0.05 vs. sham and #p<0.05 vs. vehicle-treated mice). IL, interleukin; MIP-2, macrophage inflammatory protein-2.
