## Supplemental Table for "Extracellular CIRP as a Novel Endogenous TREM-1 Ligand to Fuel Inflammation"

**Supplemental Table 1:** Primer sequences for RTqPCR.

| Gene | Ref Seq | Forward primer | Reverse primer |
|---|---|---|---|
| β-actin | NM_007393 | CGTGAAAAGATGACCCAGATCA | TGGTACGACCAGAGGCATACAG |
| TREM-1 | NM_021406 | CTACAACCCGATCCCTACCC | AAACCAGGCTCTTGCTGAGA |
| IL-6 | NM_031168 | CCGGAGAGGAGACTTCACAG | CAGAATTGCCATTGCACAAC |
| IL-1β | NM_008361 | CAGGATGAGGACATGAGCACC | CTCTGCAGACT-CAAACTCCAC |
| KC | NM_008176 | GCTGGGATTCAC- CTCAAGAA | ACAGGTGCCATCAGAGCAGT |
| MIP-2 | NM_009140 | CCCTGGTTCAGAAAATCATCCA | GCTCCTC- CTTTCCAGGTCAGT |
| TNF-α | NM_012675 | AGACCCTCACACTCAGATCATCTTC | TTG CTACGACGTGGGCTACA |
| ICAM-1 | NM_010493 | GGGCTGGCATTGTTCTCTAA | CTTCAGAGGCAGGAAACAGG |
| VCAM-1 | NM_011693 | GAACCCAAACAGAGGCAGAG | TGAGCAGCTCAGGTTCACAG |
